# Frequent female song in blue tits: behavioural context suggests a role in intrasexual competition

**DOI:** 10.1101/2021.07.01.450672

**Authors:** Javier Sierro, Selvino R. de Kort, Katharina Riebel, Ian R. Hartley

## Abstract

The blue tit (*Cyanistes caeruleus*) is an important avian model in evolutionary ecology (> 20,000 published scientific studies). Song, like in other songbird species, is generally described as a male trait and plays an important role in mate attraction and territory defence. Over the decades, there have been recurring reports of anecdotal female song but these have not led to any quantitative study of female song in blue tits. Our systematic sampling over three years revealed prolific female singing in a northern population of colour ringed blue tits. Daytime singing of females occurred throughout the breeding season during agonistic interactions, solo songs and alarm situations, and these contexts are similar to male song. Notably, female song was absent during the dawn chorus; the period around sunrise when males sing intensively just before mating. Female and male song overlapped substantially in acoustic structure (i.e. same song types, peak frequency or trill rates) but there were also significant differences in that females had smaller repertoires, shorter trills and lower vocal consistency. Differential selections pressures related with contextual (functional) differences in the role of male and female song could explain the observed differences in acoustic structure. The new finding of prolific female singing in such a well-studied species suggests we ought to revise our understanding of male and female vocal communication in this (and probably other) species. Identifying the selection pressures associated to the convergence versus divergence of male and female song may provide important insight in understanding birdsong evolution.

**Lay summary:** Female song has been anecdotally reported in blue tits but there are no quantitative studies of singing behaviour, song structure or context. Here, we report frequent female singing in blue tits, associated mostly with agonistic interactions and alarm situations. Importantly, female song was not observed during dawn chorus, the period around sunrise when males sing intensively just before mating. Female and male song overlapped substantially in acoustic structure (i.e. same song types) but there were also significant differences (i.e. females sang with lower vocal consistency). We speculate that differences in context (function) of male and female song could explain the observed differences in acoustic structure. The new finding of prolific female singing in such a well-studied species suggests we ought to revise our understanding of male and female vocal communication.

## INTRODUCTION

Birdsong plays an important role in the acquisition of breeding resources, to mediate social conflicts and to attract mates (Catchpole and Slater 2008; Marler and Slabbekoorn 2004) but also for pair coordination and in alarm situations (Cresswell 1994; Halkin 1997). Birdsong is, therefore, under strong sexual selection (but see Tobias et al. 2011) and it has been assumed to be, predominantly, a male trait (Collins 2004; Searcy and Andersson 1986; Searcy and Yasukawa 1990). Consequently, its function has been mostly studied in males (Austin et al. 2021; Langmore et al. 1996; Odom et al. 2014; Riebel et al. 2005) regardless of many reports of female song right from the early days of modern birdsong research (e.g. Nice 1943; Robinson 1949, Hinde 1952; Hoelzel 1986; Ritchison 1986).

It was not until this century that the first systematic worldwide survey was conducted showing that female song is common, particularly in the basal clades of passerines, making concurrent male and female song the most likely ancestral state (Odom et al. 2014). With this shifting view, new questions arise regarding the function of female song and the selection pressures underlying sexual differences. Even though the study of female song lacks the literature background of male song, the evidence indicates that it can serve similar functions such as territory advertisement (Cain et al. 2015; Cooney and Cockburn 1995), mate attraction (Langmore et al. 1996), mate guarding (Reichard et al. 2018) or resource defence (Tobias and Seddon 2009). One of the most common roles of female song, regarding non-duetting species, is related to the competition for breeding resources (and mates) between females (Austin et al. 2021; Langmore 1998). Systematic research is needed to gain a complete picture of shared versus sex-specific functions of song in passerines (Austin et al. 2021; Riebel et al. 2019).

One of the unresolved issues is why female song is common in the (sub)tropics and in ancestral clades, while it seems rare in temperate zones and in Passerida. A current working hypotheses is that short breeding seasons, seasonal territoriality and migration might be associated with the loss of song (Benedict 2008; Odom et al. 2014; Price 2009). However, the rare documented cases of functional song in Passerida could have partly arisen from sampling biases (sexing a singing bird as male) (Odom and Benedict 2018). Furthermore, there have been especially few systematic studies in Northern Temperate regions to quantify female song and its function(s) (Riebel et al. 2019). Important first steps are the documentation of female song across and within species, finding ecological correlates, describing behavioural contexts and sex specific structure of song.

The blue tit might be a case in point: this common, widespread and non-migratory passerine breeds in the temperate regions of Europe and western Asia. It is a model species for studies of birdsong, mating systems and other aspects of behavioural ecology (reviewed in Mainwaring and Hartley 2019), with more than 20,000 scientific publications. To human observers, males and females show only minor sexual plumage and size dimorphism and there is much overlap in colour intensity and size between the sexes. While there are recurring anecdotal reports of female song (Bijnens and Dhondt 1984; Cramp and Perrins 1993; Hinde 1952; Mahr et al. 2016), there are no detailed, quantitative descriptions of female singing behaviour, song structure or context, despite the extensive literature of song research in this species (Doutrelant et al. 1998; Doutrelant et al. 2000a; Doutrelant et al. 1999; Doutrelant et al. 2000b; Gorissen and Eens 2005; Gorissen et al. 2002; Hinde 1952; Latimer 1977; Poesel and Dabelsteen 2005; Poesel et al. 2004; Poesel et al. 2001; Poesel and Kempenaers 2000; Stadler 1951)

Overall, from the literature one gains the impression that female song is rare in blue tits. In contrast, during a systematic song recording effort in an individually colour-ringed population, we encountered frequent female song throughout the breeding season. Here, we present a quantitative analysis of the context, occurrence and acoustic structure of female song, in comparison with the song of their male partners. Blue tits generally breed in monogamous pairs, but are occasionally socially polygynous and frequently genetically polyandrous (Leech et al. 2001; Schlicht and Kempenaers 2021). In other facultative polygynous species, females use song as a mate-guarding signal during female-female competition (Austin et al. 2021; Langmore 1998). If this is the same for blue tits, we would predict that females should sing during agonistic interaction with other females. If the function of female song in blue tits is associated with territory defence, we would expect to find females producing solo songs, as a territory advertisement, and during agonistic interactions with both sexes. Solo singing by females would also be consistent with a potential function in mate attraction. Furthermore, female song during the dawn chorus, a time when copulations take place in blue tits, could also be interpreted as providing a function in mate attraction or seeking within- or extra-pair copulations. Finally, we discuss the sexual similarities and differences in song structure in relation to possible functional differences based on the context where we find females and males singing. Results derived from this study might provide clues with respect to potential sex-specific functions of female song and help with developing testable hypothesis for future experimental work.

## METHODS

### Study species and sampling methods

Blue tit song is usually composed of a few introductory, high-pitched notes followed by a trill, defined as the last part of the song where a note is repeated in succession (Figure 1), (Bijnens and Dhondt 1984; Cramp and Perrins 1993). A note was defined as a continuous trace in the spectrogram, separated from other notes by silent gaps. Each individual presents several stereotyped song structures referred to as song types (see also ‘phrase’ in Poesel and Kempenaers 2000) and these are usually comparable between individuals within the population (Figure 1). During singing, blue tits repeat the same song type many times, alternated with silent pauses (i.e. discontinuous singers), before switching to a different song type which results in so-called song type bouts.

**Figure 1.**
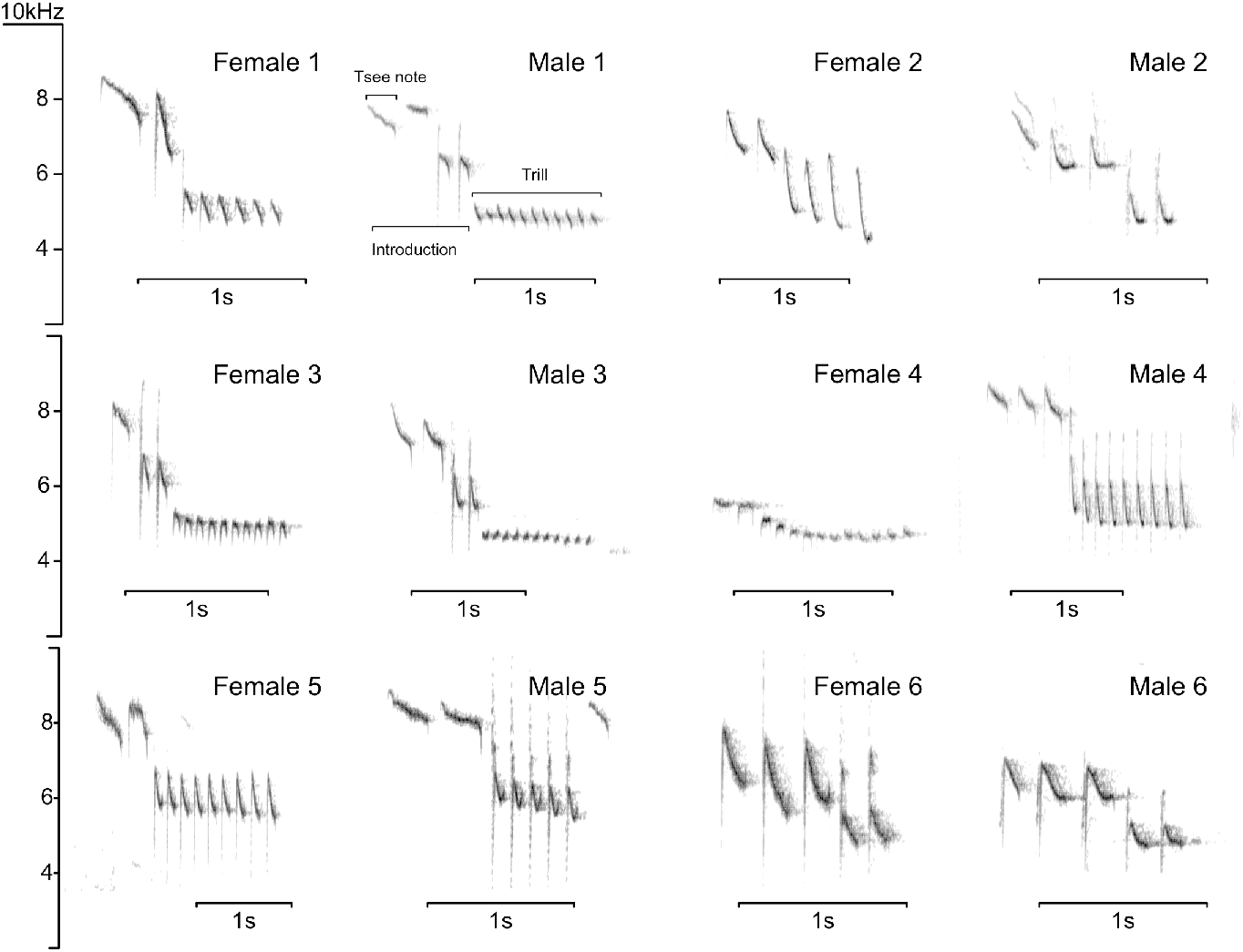
Spectrograms of female and male songs. For each female, we selected the same, or similar, song type of her mate for display. The basic features of blue tit song structure are indicated in the first example. The songs were selected to provide a good visualization of song structure, not necessarily a statistical representation of sexual differences.

All birds observed in this study were part of a breeding population occupying nest boxes in deciduous and mixed woodland at Lancaster University campus (54.01° N, 2.78° W) and form part of long-term monitoring study based on approximately 120 available nest boxes (Mainwaring and Hartley 2009). Each year, adult breeding pairs are caught, measured (straightened, flattened, wing length to nearest mm, tarsus length with foot bent down, to nearest 0.1 mm and head-bill length to nearest 0.1 mm), weighed (to nearest 0.1 g) and ringed with a unique combination of three coloured rings and one numbered metal ring (Redfern and Clark 2001). During the breeding season, individuals were sexed in the hand based on the presence of a brood patch or cloacal protuberance (Svensson 1992). Birds were aged as first-year or older than first-year, based on plumage characteristics described in (Svensson 1992).

From Jan-May 2018 to 2020, we conducted daily walked transects to collect song recordings using a Marantz PMD661 recorder (48kHz sampling rate and 24-bit depth) and a Sennheiser ME67 microphone. We followed linear transects since nest boxes were placed in lines within strips of woodland (Figure S1). The blue tits found along the transect were identified from their colour rings. The study was initially designed to investigate variation in song performance of males, but when collecting male recordings, we regularly encountered singing females and decided to concurrently establish a data base of female song (i.e. Wilkins et al. 2020). When a blue tit was encountered singing, its behaviour and song were recorded simultaneously. The observation ended whenever the focal bird stopped singing for more than a few minutes or went out of sight. In the field, the sex of the singing bird was generally unknown to the observer, but using the ring details recorded, individuals could be sexed after cross-checking with the database. We used a dictaphone to take voice notes in a continuous recording for each day of fieldwork. The dictaphone also recorded the ambient sounds, including songs of blue tits that were nearby and this was useful to record the first songs of a bout if song was unpredictable, just before we started the high-quality audio recorder. During the pre-breeding period, from January to the beginning of April, we collected song during daytime singing, after sunrise throughout the morning since singing activity was still very low at dawn (Hinde 1952). In April and May, during the egg-laying (fertile) and incubation periods, when dawn singing behaviour was more predictable (Hinde 1952), we began sampling one hour before sunrise until one hour after sunrise. At dawn, visibility was poor and therefore identification of birds using colour rings was difficult. To confirm the identity of a bird singing at dawn, we kept following and recording it until light levels were high enough for identification.

### Categorizing the behavioural context of blue tit song

After a first revision of song recordings, we chose four distinct behavioural contexts to categorize all singing instances: alarm, agonistic interactions, solo song and dawn song (see Table 1 for operational definitions). For each bird we categorized every singing instance recorded into one of the four behavioural contexts and then estimated the proportion of each context for each individual. We then selected a set of males that were partners of the recorded females and conducted a similar categorization of behavioural context for all singing instances recorded.

**Table 1.** Operational definition of behavioural and acoustic terms

| Term | Definition |
| --- | --- |
| Alarm display | Characterised by the production of 1) isolated, high-pitched ‘tsee’ calls or scolding calls, both typically produced in the presence of a predator (Bijnens and Dhondt 1984; Hinde 1952; Latimer 1977; Mahr et al. 2016) or 2) mobbing behaviour such as approaching closely and circling a potential predator (Cramp and Perrins 1993; Shirihaï and Svensson 2018). |
| Solo songs | Characterised by an individual bird singing alone without any active social interactions with conspecific or alarm responses. |
| Agonistic interactions | Defined as any interaction where two or more blue tits actively chased, displaced or attacked each other (Irschick et al. 2007). In some cases, the identity and therefore sex of the non-focal individual(s) was unknown and then we did not further classify these interactions into intra- or inter- sexual interactions. |
| Dawn song | A long vocal display that starts 90-30 min before sunrise, in line with literature on this and other species (Gil and Llusia 2020; McGregor 2005; Poesel et al. 2001) |
| Song repertoire | Total number of song types found across all recordings of a single individual |
| Peak frequency | The maximum amplitude frequency in the mean power-spectrum (window size = 2048, window type = ‘hanning’) |
| Trill length | Duration of the trill in seconds |
| Trill rate | Number of notes per second always excluding the last rendition of the trill to avoid bias of higher rates toward shorter trills |
| Vocal deviation | Using song of both sexes, we first calculated the upper bound regression between trill bandwidth and trill rate using bins of 100 Hz (Blackburn et al. 1992; Podos 1997). The vocal deviation is the orthogonal distance of each trill to the estimated upper bound limit (Figure 4, see Podos et al., 1997). |
| Vocal consistency | Acoustic similarity between each note to the consecutive rendition within the trill using a spectrogram cross-correlation algorithm (SPCC) (Botero et al. 2009; Clark et al. 1987; Coleman et al. 2007). We calculated the maximum correlation of every pairwise SPCC with a maximum temporal offset of 20 ms and a temporal resolution of 1 ms. The spectrogram matrices were computed using an FFT algorithm with a window size of 512 samples and 90 % overlap between successive windows and ‘Hanning’ window type. |

### Song type repertoire analysis

Blue tits have relatively small individual repertoires, ranging from 3-8 different song types (Bijnens and Dhondt 1984). Since blue tits repeat the same song many times before switching, repertoire size is often estimated on the basis of the long, sustained singing displayed during dawn song (Doutrelant et al. 2000a; Poesel et al. 2004). We did not record cases of females singing dawn song, and female daytime singing, like that of males, consisted of much shorter singing bouts. Hence, to compare repertoire usage between males and females, we focussed on daytime singing and counted the number of distinct song types across several recording days of the same individual. Following this method, we selected all those females recorded on two or more dates, and called this the repertoire data set. We categorized song types by visual inspection of spectrograms (Bijnens and Dhondt 1984; Doutrelant et al. 1998) in Audacity (Mazzoni and Dannenberg 2014) (window type: ‘Hanning’, window length 1024 samples, 90% overlap and -80 dB range). Based on the song delivery mode of blue tits, that repeat the same song type many times, we focused on the switching points between song types within individuals as the main criteria for identifying song types. The switching point between song types was easily identifiable even if song types were similar. Some of the main features of song that helped in categorizing song types were trill rate, frequency modulation of trill notes, the trill length and the structure of the introductory part (Figure 1). For every female, we selected her male partner to compare song repertoires between sexes. For all pairs, we collected a larger recording sample for the male than for the female for both the number of songs and recording dates. We used a random number generator to select, for each paired male, exactly the same number of songs as his female partner, recorded on the same number of dates choosing the nearest dates in relation to the fertile period (day of first egg) and excluding dawn chorus recordings. While this means that we might not have included the complete repertoire of each individual, this sampling method allowed us to compare repertoire usage between the sexes.

### Acoustic analysis

For detailed analysis of acoustic parameters of female song, we chose a subset of females with song recordings of high signal-to-noise ratio, hereafter the acoustic data set (Table S1). We then selected individual males that were breeding partners of these females in at least one breeding season. Again, for each male partner, we selected a random sample of songs, recorded in similar days in the season and excluding songs recorded during the dawn chorus (Table S1).

We conducted all acoustic analysis in R (*tuneR*: Ligges 2013, *seewave*: Sueur et al. 2006; R Development Core Team 2019). For each individual, we analysed a maximum of ten songs for each song type and each date recorded. Acoustic measurements were made in the terminal trill only since the introductory notes of blue tit songs are more variable and may be absent in some song types (Figure 1). Despite having selected high quality recordings, we still had to exclude some notes that were masked by extraneous sounds but included the rest of the song in the analysis (7.04 ± 5.3% of notes in females and 6.4 ± 5.5% of notes in males). We did this to avoid biasing the sample towards shorter songs, as longer songs were more likely to be partly masked.

Every note was manually labelled in the spectrogram, using the cursor to mark the start and end times. Each note was then cut out of the recording and saved as a single, normalized *wav* file. Following Podos (1997), we measured the peak, maximum and minimum frequency of each note in the mean power-spectrum (window size: 1024 samples; amplitude threshold; -20 dB) and from this we derived the bandwidth of the note. Apart from spectral features, we also took measurements of song performance including vocal consistency (*sensu* de Kort et al. 2009), trill length, trill rate and vocal deviation (*sensu* Podos 1997). Table 1 shows detailed operational definitions of each acoustic variable. All acoustic variables were measured in each note and summarised as mean values per song unit for statistical analysis.

### Statistical analysis

We created three data sets from the same group of females, one to investigate behavioural context of song, one to investigate song type repertoire and one to investigate acoustic structure in female and male song. The same individuals could be part of one or all three data sets. For a summary of the three data sets for each analysis see Table S1. All measures are presented as mean ± one standard deviation (SD), unless otherwise indicated. Statistical analyses were carried out in R software (R Development Core Team 2016).

We explored the sexual differences in singing context by comparing the proportion of singing instances in each behavioural context within individual, excluding the singing instances of males during the dawn chorus. For this we fitted Linear Mixed-effects Models (LMM) with the proportion of singing instances of each individual as a function of sex and its interaction with the behavioural context (Table 2). To account for repeated measurements we included the individual identity as a random effect. We also included the pair identification as a random effect to group together male and female breeding partners. In the behavioural data set, we had different numbers of males and females because some males were not recorded and, in other cases, different females had the same partner in different years (see *Results*). Using LMM allowed us to cope with this unbalanced design but, in any case, all males included in the analysis were partners of at least one of the females. To investigate changes in song production through the season we estimated the Pearson’s correlation coefficient between the proportion of singing females, per female identified, with the week in relation to the first egg date.

**Table 2.** Model output comparing behavioural context of song between sexes. For each fixed effect we present the model estimate, the lower and higher CI of the estimate, the Z statistic derived from Wald tests and the relative importance of that factor in the final model. The last two columns show the marginal R squared represented as R^2^_m_ and the conditional R-squared represented as R^2^_c_ of the full model. This output shows that proportion of song observations in solo songs, agonistic context is similar between sexes, whereas in the alarm context, females were observed significantly more often in proportion.

| Parameter | Fixed effect | Estimate | 2.5 % CI | 97.5 % CI | Z value | Relative importance | $R^2_m$ | $R^2_c$ |
| --- | --- | --- | --- | --- | --- | --- | --- | --- |
| Behavioural context of song | Intercept | 0.73 | 11.564 | 0.606 | 0.853 | - |  |  |
|  | Context (Alarm) | -0.007 | 0.063 | -0.214 | 0.201 | 1 | 0.246 | 0.495 |
|  | Context (Agonistic) | -0.151 | 1.752 | -0.321 | 0.018 | 0.766 |  |  |
|  | Sex (male) | -0.064 | 0.695 | -0.243 | 0.116 | 0.958 |  |  |
|  | Context (Alarm) : Sex (male) | -0.38 | 3.609 | -0.586 | -0.174 | 0.766 |  |  |
|  | Context (Agonistic) : Sex (male) | -0.16 | 1.505 | -0.367 | 0.048 | 0.958 |  |  |

For the song repertoire analysis we compared females and males for the number of song types within the sample of each individual using Wilcoxon signed-rank tests.

For the acoustic analysis, we built five LMMs to test for sex differences for each of the following five parameters: peak frequency, vocal consistency, trill length, trill rate and vocal deviation (Table 3). To define the spectral features of song, we selected only the mean peak frequency of each song, because it was the most robust measurement and it was strongly correlated with the maximum frequency (*r* (798) = 0.90, *P* < 0.001) and the minimum frequency (*r* (798) = 0. 89, *P* < 0.001). As explanatory variables we used sex (male, female), age (first-year or older than first-year) and weeks in relation to first egg date to account for seasonal variation (week of first egg = 0) (*sensu* Schlicht and Kempenaers 2020). If breeding data were missing for a particular individual, we used the mean date of the first egg in our study site for that year to estimate weeks to first egg. We included sex-specific interactions with age and season effects (in weeks in relation to first egg) to investigate their potential effects on each sex independently. To model peak frequency, we also included the tarsus length and its interaction with sex, since this song feature could be affected by sexual dimorphism in body size. To account for repeated measurements, and to match female and male paired partners, we included the individual identity and the pair identification as random effects in all models.

**Table 3.** Model output of the full average model for each song trait, comparing male and female song. For each fixed effect we present the model estimate, its corresponding standard error, the lower and higher CI of the estimate, the Z statistic derived from Wald tests and the relative importance of that factor in the final model. The last two columns show the marginal R squared represented as R^2^_m_ and the conditional R-squared represented as R^2^_c_ of the full model. Overall, we found a significant difference between sexes in vocal consistency and trill length.

| Parameter | Fixed effect | Estimate | 2.5 % CI | 97.5 % CI | Z value | Relative importance | $R^2_m$ | $R^2_c$ |
| --- | --- | --- | --- | --- | --- | --- | --- | --- |
| Peak frequency (kHz) | Intercept | 5.073 | 4.956 | 5.191 | 84.541 | - |  |  |
|  | Weeks to first egg | -0.136 | -0.199 | -0.072 | 4.191 | 1 | 0.08 | 0.644 |
|  | Age | -0.055 | -0.101 | -0.009 | 2.328 | 0.201 |  |  |
|  | Tarsus length | -0.096 | -0.194 | 0.001 | 1.93 | 0.175 |  |  |
| Trill length (s) | Intercept | 0.451 | 0.367 | 0.535 | 10.531 | - | 0.126 | 0.604 |
|  | Sex (male) | 0.224 | 0.103 | 0.347 | 3.609 | 1 |  |  |
| Vocal deviation (orthogonal distance) | Intercept | 6.058 | 5.244 | 6.871 | 14.59 | - | 0.017 | 0.286 |
|  | Sex (male) | 0.702 | -0.514 | 1.918 | 1.132 | 0.517 |  |  |
|  | Age (older) | 0.303 | -0.1 | 0.706 | 1.475 | 0.357 |  |  |
| Vocal consistency (SPCC score) | Intercept | 0.869 | 0.841 | 0.898 | 59.574 | - | 0.125 | 0.416 |
|  | Weeks to first egg | 0.038 | 0.024 | 0.051 | 5.541 | 1 |  |  |
|  | Sex (male) | 0.042 | 0.013 | 0.071 | 2.828 | 0.37 |  |  |
| Trill rate (notes per sec) | Intercept | 8.52 | 7.926 | 9.114 | 28.115 | - | 0.04 | 0.732 |
|  | Sex (male) | -0.125 | -1.163 | 0.913 | 0.236 | 0.33 |  |  |

To validate all models, we confirmed that the residuals were homoscedastic and close to a normal distribution using diagnostic plots (Knief and Forstmeier 2021; Zuur et al. 2009). To find which factors were important in explaining variation in song we computed all possible model combinations ranking them by the Akaike Information Criterion for small samples (AICc). This procedure compares the fit of all possible models while penalizing the complexity, in terms of the number of explanatory variables included. We selected all models that had ΔAICc < 2, in relation to the model with the lowest AICc score, to compute the full average model as the final model (Burnham and Anderson 2002; Burnham et al. 2011). We used the relative importance of each factor in the final model together with the coefficients and estimated confident intervals (CI) with a threshold of 95% (Burnham et al. 2011; Nakagawa and Cuthill 2007), considering there was a significant effect if the CI did not overlap with zero. We considered there was a non-significant trend when the 90% CI did not overlap with zero, and in those cases, it is embedded in the text. Finally, we calculated the *R*^2^ _GLMM_ of the full models to measure the goodness of fit (Nakagawa and Schielzeth 2013). All numerical variables are scaled and centred so model estimates are standardized (Gelman 2008). All the model coefficients are reported in Tables 2 & 3.

## RESULTS

### Behavioural context and singing activity

We observed a female singing every 2.65 hours in the field, and these were 36 ± 42 % of all females identified per day. For the same period, we recorded a male singing every 1.26 h of fieldwork, and these were 53 ± 44 % of all males identified per day. Females were observed and recorded singing during the entire study period from January to the beginning of May and we did not find any significant change in the proportion of singing females through the season (*r*(16) = -0.22, *P* = 0.38) but we did not record any singing females in the last two weeks of May (mean first egg date during the three years was the 22^nd^ of April).

In total, we gathered 99 instances of singing from 36 different females and 192 singing instances from 31 selected male partners (Table S1). From all observations of female singing, 42% were during alarm displays, 42% in agonistic interactions and 16% during spontaneous song (Figure 2). During agonistic interactions, we recorded two cases where females produced song along with direct physical aggression directed towards other females. No female was observed singing during the dawn chorus (i.e. concurrent with the long singing displays, starting 90-30 min before sunrise (Gil and Llusia 2020; McGregor 2005; Poesel et al. 2001). In males, 34% of the total of 192 singing instances were observed during the dawn chorus. After excluding dawn chorus observations, we were left with 127 observations, 61% during spontaneous song, 19% during alarm displays and 20% during agonistic interactions. Our results show that, in proportion, females use song in alarm behaviour significantly more often than males (Table 2, Figure 2). On the other hand, the proportions of song during agonistic interactions and spontaneous song were not significantly different between the sexes (Table 2, Figure 2).

**Figure 2.**
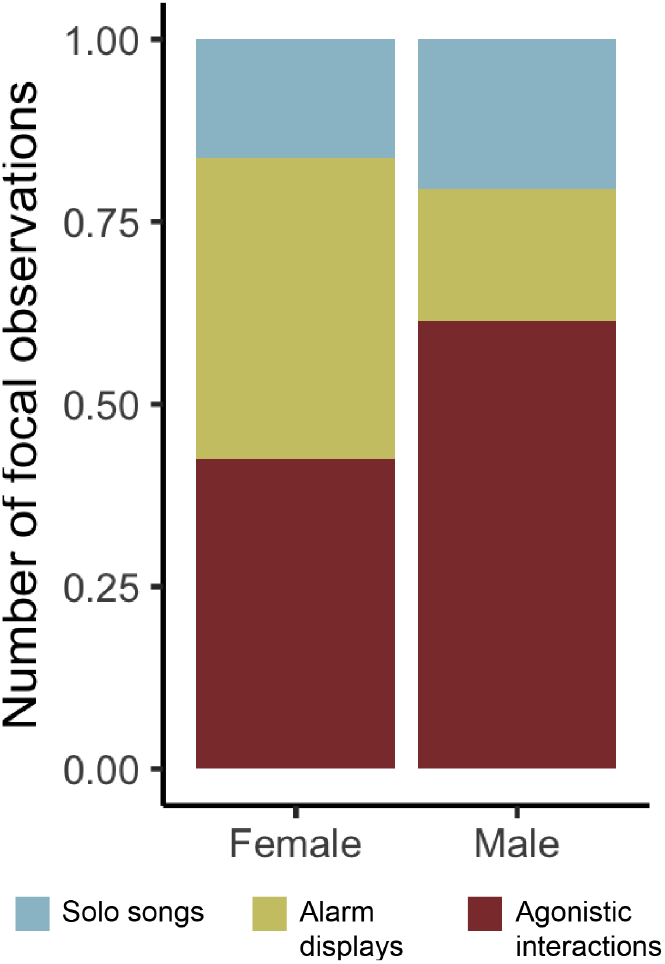
Behavioural context of song production in females (N = 99 observations) and males (N = 127 observations), after removing observations of males during dawn chorus. Results indicate there is a significant difference in the proportion of focal observations in alarm displays between sexes, but not in other contexts.

### Song repertoire

Song repertoires were assessed for 19 females that were recorded at least on two different dates (60.5 ± 43.5 songs and 3.8 ± 1.8 dates per female) and their 19 male partners (58.4 ± 40.2 songs and 3.8 ± 1.8 dates per male, Table S1). Following visual inspection of songs, we found that males and females used the same song type categories but the number of song types per individual sample was significantly lower in females than in males (Females = 1.50 ± 0.70 versus males = 2.65 ± 0.96 song types; N=19 pairs, W = 63, *P* < 0.001, 5% CI = -1.00, 95% CI= -2.00; Figure 3). For 13 of 19 females we only recorded one song type, in contrast, none of males had fewer than two song types in this or other populations (Bijnens and Dhondt 1984; Poesel et al. 2001). Three of the most recorded females sang only one song type, with more than 5 dates of recording, in two (or more) years and approximately 100 songs sampled, (Table S1).

**Figure 3.**
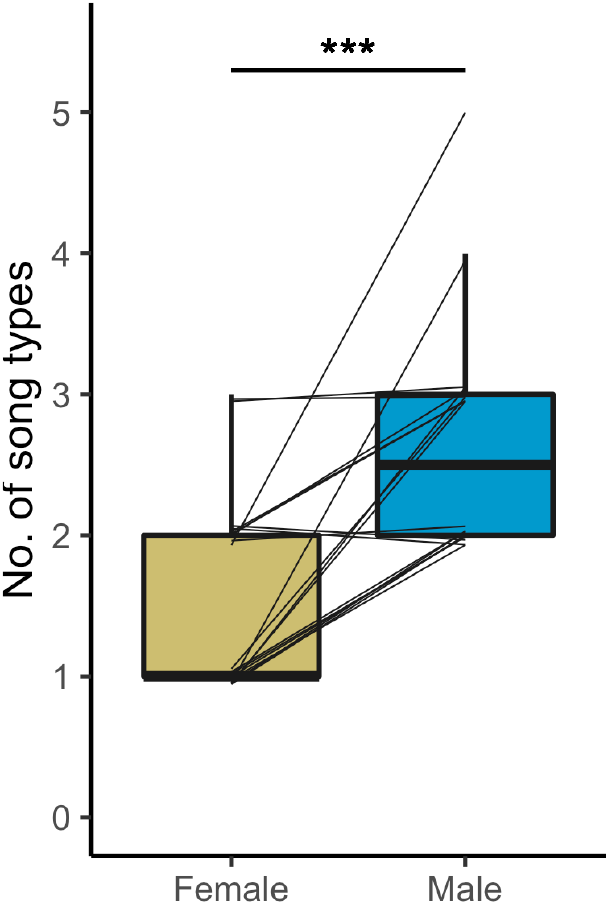
Comparative analysis of song repertoire between sexes, measured as number of song types per individual across all recordings collected during the sampling period (only females recorded at least in two dates). Females are shown in gold and males in blue with lines connecting female-male partners. *** = *P* < 0.001, Wilcox Signed-Rank test (Females = 1.50 ± 0.70 versus males = 2.65 ± 0.96 song types; N=19 pairs, W = 63, *P* < 0.001, 5% CI = -1.00, 95% CI= -2.00).

### Acoustic analysis: Spectral and performance variables

For the description of spectral and performance features of female song and subsequent comparison with the songs of their breeding partners, we selected 402 high audio quality songs from 32 females (12.6 ± 12.3, songs per individual) and 345 high audio quality songs from 28 male partners (12.3 ± 12.2, songs per individual, Table S1). Female blue tits sang with significantly lower vocal consistency and produced shorter trills than males (Figure 4, Table 3), but they did not differ in peak frequency, trill rate or vocal deviation (Figure 4, Table 3). Upper bound regression of vocal deviation including male and female song rendered a significantly negative slope (*r*(35) = -0.90, *P* < 0.001, Figure S2). In the case of vocal consistency, both males and females increased vocal consistency during winter towards spring (fertile period), independently of age and sex (Figure 5, Table 3). In both males and females, peak frequency of song decreased significantly during winter towards the spring (fertile period) and older birds showed significantly lower peak frequency than first-year birds (Figure 5, Table 3). Peak frequency also showed a marginally non-significant, negative trend in relation to tarsus length in both sexes (5% CI = -0.19, 90% CI = -0.016). Note that the effects of age, season and tarsus length were estimated separately for males and females.

**Figure 4.**
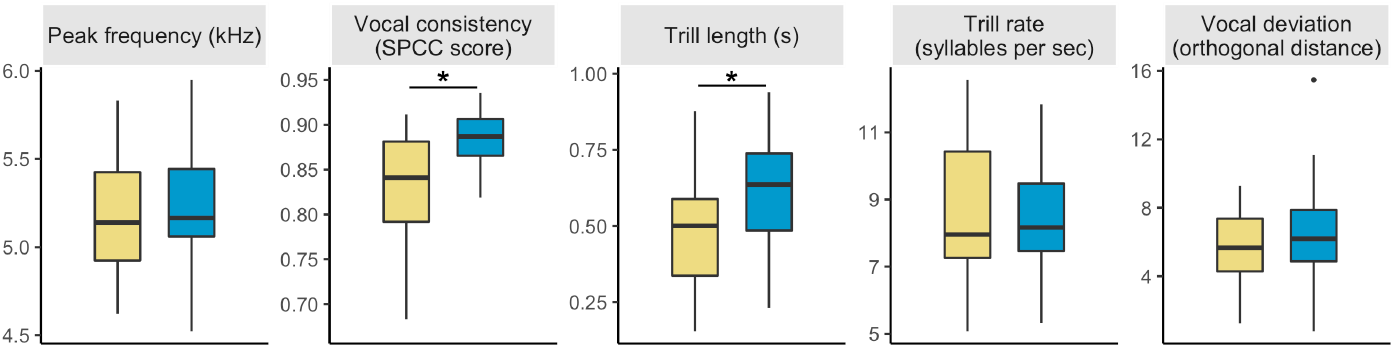
Results of sex differences in acoustic variables. Females sing with lower vocal consistency and shorter trill length than males. Peak frequency, trill rate and vocal deviation are not statistically different between sexes. Significance code: * means that the CI for the sex effect in the model does not overlap with zero.

**Figure 5.**
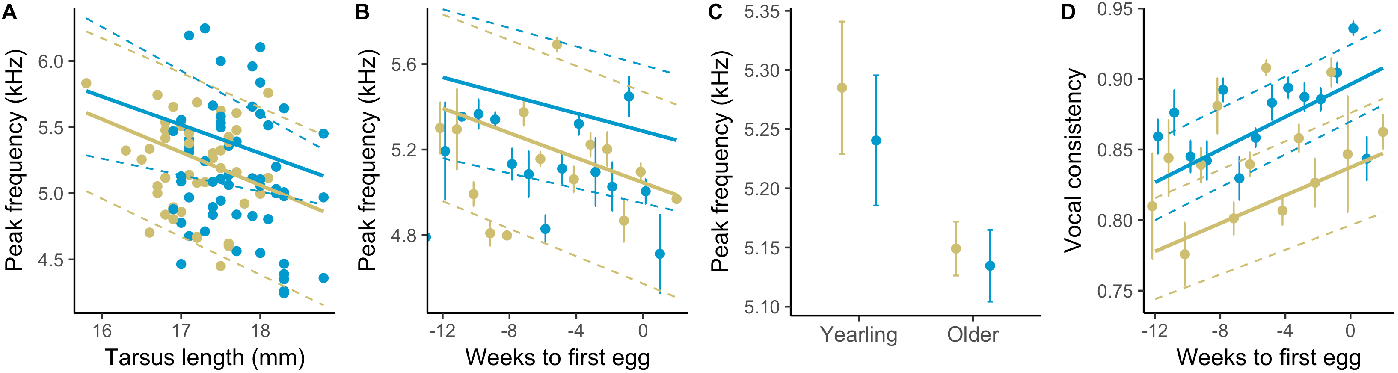
Scatter plots showing the significant effects of tarsus length, age and season on various song parameters. These effects are estimated separately and specifically for males and females in the model. Points represent raw data and lines represent predicted values from the model, with the associated confidence interval in dashed lines. Females are represented in gold and males in blue.

## DISCUSSION

In blue tits, female song was common across a variety of behavioural contexts. This contrasts with previous anecdotal reports of female song in this species, associated mostly with alarm contexts. Based on an *ad hoc* sampling scheme, registering whether each identified individual was singing during the daytime, we observed a high incidence of females singing (36%) relative to males (53%), normally considered the “singing sex”. In contrast, we did not collect any observations of females singing dawn song (but see Gorissen and Eens (2005) for female vocal activity inside the nest box). As there is little evidence that ecological or environmental factors are unusual in our population, we assume that females in other populations might sing as much. Hence, the most parsimonious explanation for the under-reporting of female song in the literature is that singing in female blue tits has been overlooked. Apart from the general research bias towards male song, two more specific reasons may account for these oversights. One, a focus of many studies on dawn song, a context when female blue tits indeed do not seem to sing, and second, the low dimorphism (to the human eye) that makes it difficult to sex birds from a distance in the field. As song is often used to sex birds, singing females might have been registered as singing males. Importantly, female song, combined with physical aggression, was observed only during female-female interactions, which is in line with a potential function in intrasexual competition, supporting previous reports of female singing and fighting (Hinde 1952; Kempenaers et al. 1995). Female, like male song, showed seasonal variation in relation to the fertile period in two key parameters: peak frequency and vocal consistency. Moreover, peak frequency was related to age and weakly to body size in both sexes, indicating that female song could potentially inform receivers about the qualities of an individual singer. By comparing male and female partners, we found that individual females sang fewer song types than males and their songs were significantly shorter and lower in vocal consistency. The three most recorded females had a repertoire of one song type and this contrasts with males, that have not been reported with fewer than two song types (Bijnens and Dhondt 1984; Poesel et al. 2001). Finally, female song was similar to male song in peak frequency, trill rate and vocal deviation, raising the question as to whether some function of song is shared in both sexes. Additional work and experimental approaches are necessary to fully characterise all functions of female song, but our observational data allow us to formulate some hypotheses for future work.

First, it is important to realise that functions of female song might partly overlap with male song, but also have different functions (Austin et al. 2021; Riebel et al. 2019). Importantly, if females sing less often than males this does not mean that female song has lower ecological importance (Austin et al. 2021). In our population, female song occurred throughout the breeding season in several behavioural contexts, most often during alarm situations. In a comparable experimental context, Mahr et al. (2016) observed at least 3 different females and two different males singing when they positioned a taxidermy mount of a sparrowhawk, *Accipiter nisus*, near nests with fledglings. Our extensive observations of song during mobbing events and predator presence support those findings but show that such behaviour occurs throughout the breeding season and not only during the fledging period. Blue tits are thus among a growing number of species known to produce song in the presence of predators (Cresswell 1994; Laiolo et al. 2004; Langmore and Mulder 1992). However, anti-predator behaviour was but one context where we observed females singing. Females, like males, were regularly observed producing solo songs and also singing during agonistic interactions. Both contexts are in line with a possible territorial function of female song (Langmore 1998) that has been demonstrated in several species using playback experiments (Cooney and Cockburn 1995; Hoelzel 1986; Krieg and Getty 2016; Magoolagan and Sharp 2018). The fact that both sexes use the same song types can also facilitate joint territory defence since song type matching is an aggressive signal in this and related species (Krebs et al. 1981; Langemann et al. 2000; Poesel and Dabelsteen 2005).

Although further empirical work is needed to establish the functional value of female song in blue tits, there are three lines of evidence from our observational study indicating that female song plays a role during intrasexual competition for breeding resources and mates in the blue tit (Clutton-Brock 2009; Langmore 1998) and supporting previous reports in the blue tit (i.e. female song in ‘reproductive fighting’ Hinde 1952). First, females sing throughout the breeding season, a time of the year when blue tits are strongly territorial. Second, female song varies seasonally in acoustic structure towards the fertile period and third, females produce song during aggressive encounters with other females. A similar role of female song during intrasexual conflicts has been described in other species. In dunnocks (*Prunella modularis*) and great reed warblers (*Acrocephalus arundinaceus*), artificially high female-female competition increased the incidence of female song during intra-sexual conflicts (Kluyver 1955; Langmore and Davies 1997). Female European starlings (*Sturnus vulgaris*) produced song during aggression directed towards caged females placed in their territory (Sandell and Smith 1997). Dark-eyed junco (*Junco hyemalis*) females sang while showing aggressive behaviour towards the presentation of caged females (Reichard et al. 2018). In barn swallows (*Hirundo rustica*) females used song to interrupt male song, potentially disrupting mate attraction by the male (Wilkins et al. 2020), and in eastern whipbirds (*Psophodes olivaceus*) females approached more closely to playback of female solo song than to male or duet songs (Rogers et al. 2007).

Many of these species, where female song has a function in intrasexual competition, are opportunistically polygynous, just like blue tits (Langmore 1998). Blue tit males try to attract secondary females after the primary female starts to lay eggs (Kempenaers 1994, 1995) and polygyny can affect female fitness, since male parental care will be reduced in their nest (Hartley unpublished data). This may explain why female blue tits show strong aggressive behaviour towards intruding females during the breeding period (Gorissen et al. 2002; Kempenaers 1994, 1995; Midamegbe et al. 2011) and we observed female song during female-female fights. Female song may, therefore, play a role in intrasexual competition over mates and territories to actively ward off polygyny. To establish whether female song in blue tits functions in mate competition (mate guarding), one possibility is to test for links between the incidence of female song and the proportion of polygynous partnerships across and within populations. Playback studies are also fundamental for testing the response of female (and male) blue tits to female song within (and outside) the breeding period.

In relation to acoustic variables of song, we found that peak frequency was related to age and (weakly) to body size in both females and males. This could play an important role during female vocal interactions as it does for female Mexican antthrushes (*Formicarius moniliger*) (Kirschel et al. 2020). Female song, like male song, varied in acoustic structure during the breeding period; decreasing in peak frequency and increasing in vocal consistency towards the fertile period. In males, such seasonal variation has been associated with specific functions of song during reproduction (Ballentine et al. 2003; Halfwerk et al. 2011). The seasonal change occurred similarly in all three years, indicating that song parameters must “return to winter values” after the breeding period (summer-autumn), completing the seasonal cycle. To our knowledge, this is the first study to document seasonal changes in acoustic structure of female song.

Female song could potentially play a role in mate choice and pair formation as females display solo song during daytime singing from winter to spring, and winter associations predict pair formation (Beck et al. 2020). But importantly, the truly qualitative difference in song between sexes was the absence of female dawn song, a song display that is associated with seeking within- and extra-pair copulations by males (Kempenaers et al. 1997; Parker et al. 2006; Poesel et al. 2004; Welling et al. 1995). Given that copulations are mostly under female control, this context implies a particular selection pressure over male song that seems absent in female song. At this point, even though we need to know more about the function of female song in blue tits, we can already deduce that selection pressures shaping this signal are not congruent between the sexes (Austin et al. 2021). Theoretically, such differential pressures could imply differences in song parameters associated with female choice during copulations. In line with this, we found that song repertoire, song length and song consistency were different between males and females. Previous studies in blue tits showed that longer songs of blue tit males are associated with higher rates of extra-pair copulations (Kempenaers et al. 1997) and a playback study on female choice showed that females had a preference for higher consistency songs in blue tits (Sierro et al., in prep). Future empirical work will have to address which sexual differences, and similarities, in song are meaningful in communication and which selection pressures favour the observed differences.

Our observations of frequent singing in female blue tits are in line with recent comparative analyses that found a higher incidence of female song in non-migratory species with low sexual dichromatism (Webb et al. 2016) that hold year round territories (Benedict 2008; Logue and Hall 2014; Odom et al. 2014; Price 2009; Riebel et al. 2019). Our comprehensive report provides an example of scarce descriptions of female song in the Northern Temperate regions (reviewed in Odom and Benedict 2018; Odom et al. 2014; Riebel et al. 2019) despite a large number of species with anecdotal reports of female song (e.g. Garamszegi et al. 2007). The behavioural contexts and seasonal variation of female song in blue tits suggest a territorial and/or a mate guarding function to avoid polygyny (Langmore 1998). Furthermore, our finding that males, but not females, produce dawn song likely implies differential selection pressures between sexes since this context is highly associated with attracting females for copulations (Austin et al. 2021). For now, our observations of prolific female singing and its associations with specific contexts provide robust foundations for further hypothesis development and testing. It is crucial to design future playback studies presenting male and female blue tits with female song, as empirical evidence is key to understand its function. Finally, increasing documentation of female song in all biogeographic regions is crucial for our understanding of the evolution of birdsong.

## SUPPLEMENTARY MATERIAL

**Table S1.**
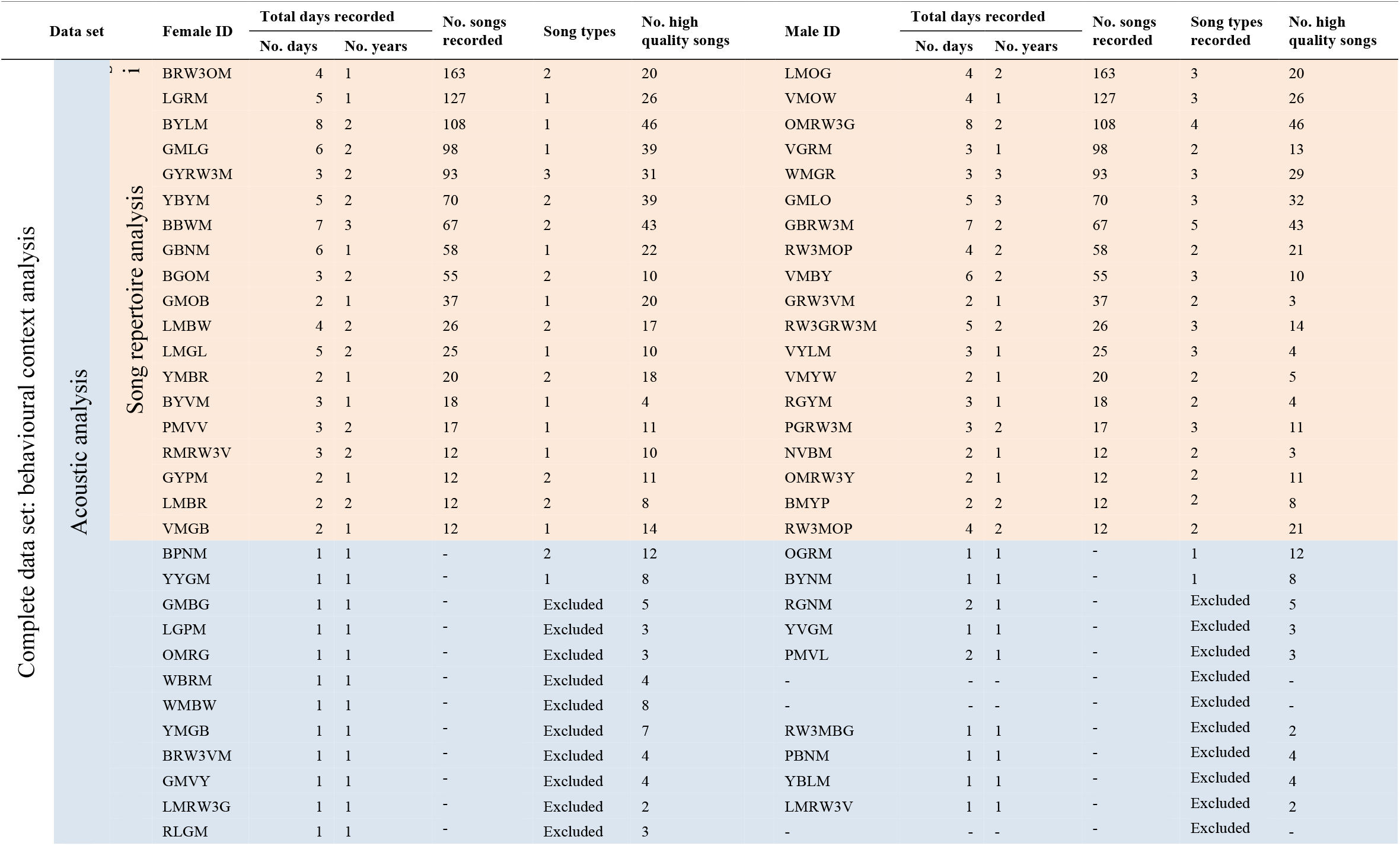
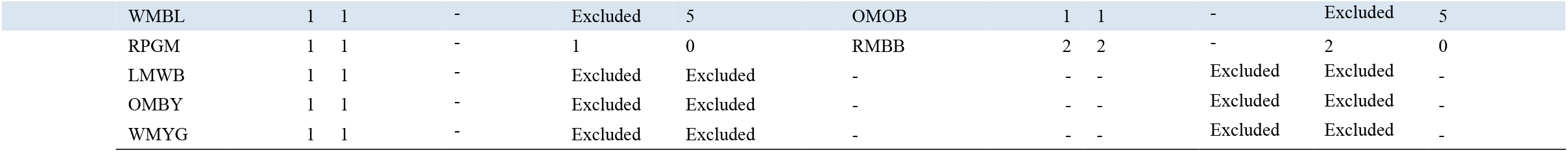
Song sample collected for each individual female, specifying the number of days and years recorded. For each female, the corresponding male partner is shown in the same row. Individuals included in the song repertoire analysis are shaded in orange. Shaded in blue (plus the orange shaded cells) are individuals included in the acoustic analysis with high-quality recordings. The entire data set represents the behavioural context analyses data set.

**Figure S1.**
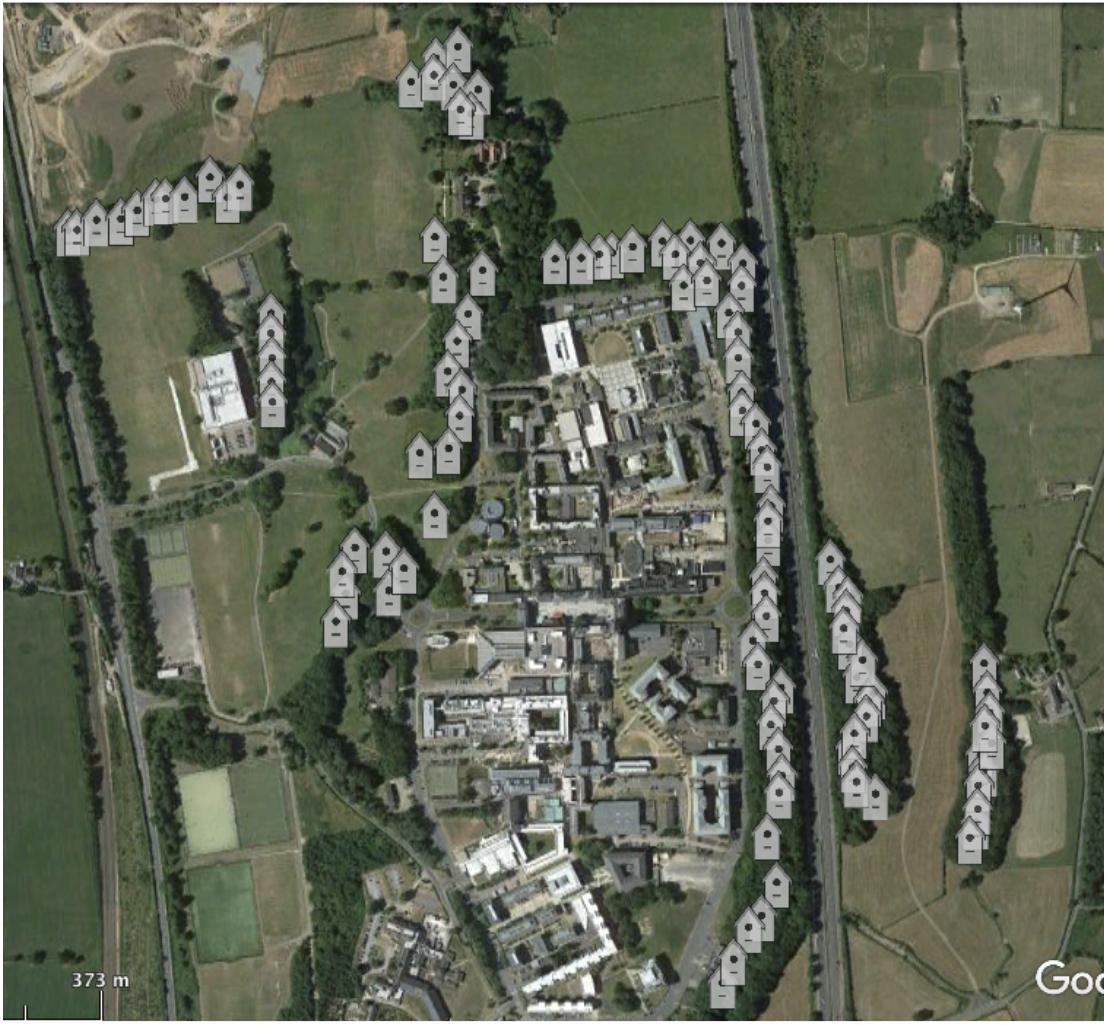
Study area where data was collected (Google Maps, 2021). The nest boxes, shown as grey symbols of a nest box, are placed in lines along the strips of forest that surround the campus of Lancaster University (54.01’ N, 2.78’ W)

**Figure S2.**
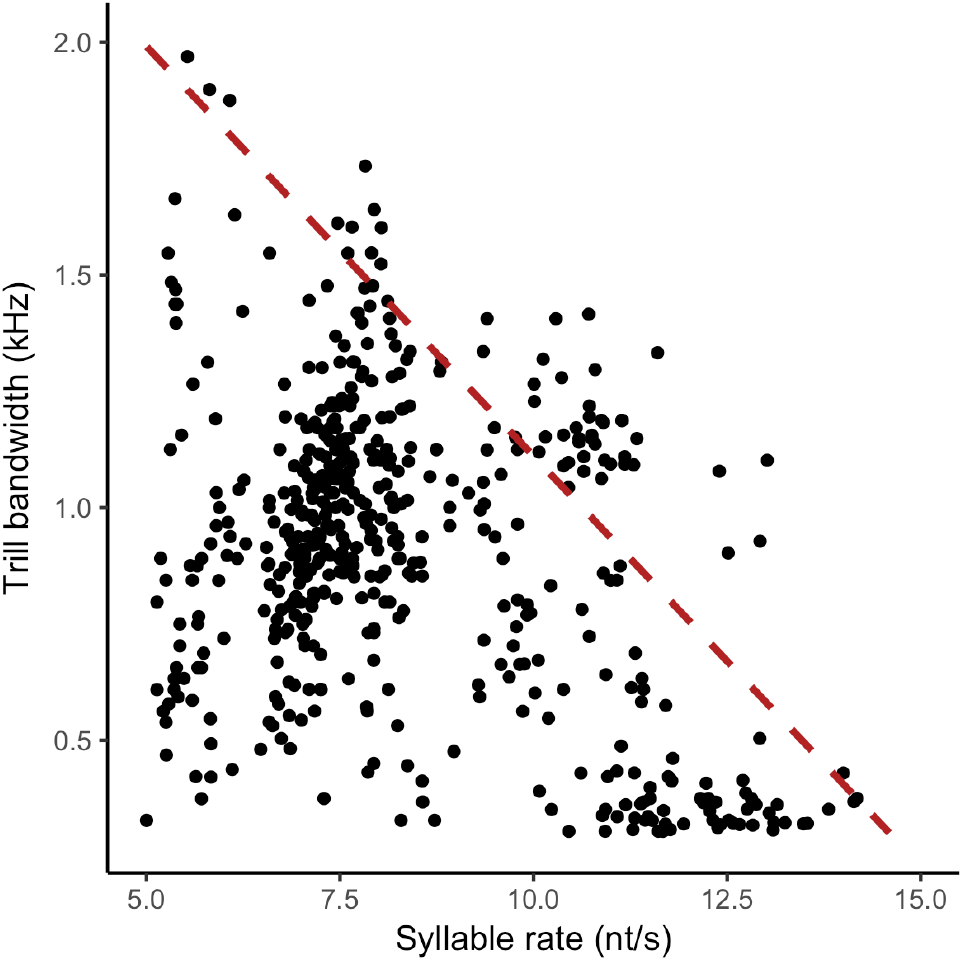
‘Triangular’ distribution of trill bandwidth in relation with trill rate, red dashed line showing the upper-bound regression (*r*(35) = -0.90, *P* < 0.001). Degrees of freedom in the upper-bound regression are related to the number of bins selected to estimate local maxima (Podos 1997). Each dot represents a song, with a total of 747 songs from males and females.

